# Integral use of immunopeptidomics and immunoinformatics for the characterization of antigen presentation and rational identification of BoLA-DR-presented peptides and epitopes

**DOI:** 10.1101/2020.12.14.422738

**Authors:** Andressa Fisch, Birkir Reynisson, Lindert Benedictus, Annalisa Nicastri, Deepali Vasoya, Ivan Morrison, Søren Buus, Beatriz Rossetti Ferreira, Isabel Kinney Ferreira de Miranda Santos, Nicola Ternette, Tim Connelley, Morten Nielsen

**Affiliations:** Ribeirão Preto College of Nursing, University of São Paulo, Av Bandeirantes 3900, Ribeirão Preto, Brazil; Department of Health Technology, Technical University of Denmark, DK-2800 Lyngby, Denmark; The Roslin Institute, Edinburgh, Midlothian EH25 9RG, UK; The Jenner Institute, Nuffield Department of Medicine, Oxford, OX37BN, UK; Laboratory of Experimental Immunology, Department of Immunology and Microbiology, Faculty of Health and Medical Sciences, University of Copenhagen, Copenhagen, Denmark; Ribeirão Preto Medical School, University of São Paulo, Av Bandeirantes 3900, Ribeirão Preto, Brazil; Instituto de Investigaciones Biotecnológicas, Universidad Nacional de San Martín, CP1650 San Martín, Argentina

## Abstract

Major histocompatibility complex (MHC) peptide binding and presentation is the most selective event defining the landscape of T cell epitopes. Consequently, understanding the diversity of MHC alleles in a given population and the parameters that define the set of ligands that can be bound and presented by each of these alleles (the immunopeptidome) has an enormous impact on our capacity to predict and manipulate the potential of protein antigens to elicit functional T cell responses. Liquid chromatography-mass spectrometry (LC-MS) analysis of MHC eluted ligands (EL data) has proven to be a powerful technique for identifying such peptidomes, and methods integrating such data for prediction of antigen presentation have reached a high level of accuracy for both MHC class I and class II. Here, we demonstrate how these techniques and prediction methods can be readily extended to the bovine leukocyte antigen class II DR locus (BoLA-DR). BoLA-DR binding motifs were characterized by EL data derived from cell lines expressing a range of DRB3 alleles prevalent in Holstein-Friesian populations. The model generated (NetBoLAIIpan - available as a web-server at www.cbs.dtu.dk/services/NetBoLAIIpan) was shown to have unprecedented predictive power to identify known BoLA-DR restricted CD4 epitopes. In summary, the results demonstrate the power of an integrated approach combining advanced MS peptidomics with immunoinformatics for characterization of the BoLA-DR antigen presentation system and provide a novel tool that can be utilised to assist in rational evaluation and selection of bovine CD4 T cell epitopes.

## Introduction

Major histocompatibility complex (MHC) genes play a vital role in the regulation of adaptive immunity. Whilst classical MHC class I genes are expressed on most nucleated cells, MHC class II (MHCII) molecules show a more restricted expression and are predominantly expressed on professional antigen-presenting cells such as dendritic cells, B-cells, and macrophages. The MHCII system enables peptides derived from both extracellular and intracellular proteins that have been delivered in the endocytic pathway to be loaded into the peptide-binding groove of MHCII molecules and be displayed as stable peptide-MHCII complexes (pMHCII) on the cell surface(1). CD4 T cells bearing cognate TCRs capable of binding specific pMHCII complexes can become activated and perform a range of functions, including supporting other immune effector cells such as macrophages, B cells and CD8 T cells(2). Thus, pMHCII molecules play a critical role in initiating and developing both humoral and cell-mediated adaptive immune responses.

MHCII molecules are heterodimers composed of an α and β chain, each consisting of an extracellular domain, a transmembrane region, and an intracytoplasmic tail. The distal membrane domains (α1 and β1, respectively) form an open peptide-binding groove that binds peptides of variable length, mainly of 13–25 amino acid residues(3). The peptide-binding groove most often contains four major pockets that interact with the side-chains of anchoring residues located at positions 1, 4, 6, and 9 of the 9-mer binding-core of the bound ligand. These pockets thus determine the binding motif of the peptides that can be presented by an MHCII molecule(4,5). A key feature of the MHC genes is the high level of polymorphism. For example in humans, three conventional MHCII heterodimers are expressed – DR, DQ and DP – and a total of ~2, ~2,500, ~100, ~1,200, ~80 and ~1,000 protein-coding variants of the α (A) and β (B) chain genes, DRA, DRB, DQA, DQB, DPA, and DPB respectively, have been identified. Except for DRA, the polymorphism of MHCII genes is focused predominantly within the α1 and β1 domains(6), resulting in variations in the residues of the binding groove, and consequently determining the variable binding motifs and so the capacity of different MHCII molecules to bind different peptide sets.

In cattle, there are only two categories of conventional MHCII molecules, BoLA-DR and BoLA-DQ(7). The DRB, DQA, and DQB genes are highly polymorphic, whilst, as in other species, the DRA gene is essentially monomorphic(8). Although there are three DRB loci, only DRB3 is considered to be functionally expressed since DRB1 is a pseudogene and DRB2 is expressed at very low levels if at all(9). Consequently, the variability of expressed BoLA-DR molecules can be characterized by sequencing of the DRB3 gene(10). The ability to perform rapid sequence-based typing of DRB3 using Sanger technology has resulted in DRB3 being the most intensely studied bovine MHC gene(11–19), with 357 alleles registered in the IPD-MHC database (November 2020: https://www.ebi.ac.uk/ipd/mhc/group/BoLA/).

Characterisation of the peptide repertoires presented by different MHCII molecules can enable the development of algorithms that predict potential MHC binding peptides within proteins rapidly. Integration of large data sets of peptides directly eluted off MHC molecules and sequenced by mass-spectrometry (MS), so-called eluted ligand (EL) data, have facilitated the generation of accurate MHC-binding prediction algorithms(20–27). Such *in silico* tools can accelerate antigen selection for vaccine development and are of particular relevance to vaccines against pathogens with large proteomes (e.g. eukaryotic parasites), where screening and selection of candidate antigens from a large number of expressed proteins would be a major obstacle.

Analysis and interpretation of EL data are made challenging by ambiguous ligand MHC assignment resulting from the multiple MHC molecules expressed on the surface of most cells. Several approaches have been proposed to address this, spanning from the engineering of cell lines and/or expressed MHC molecules to allow for analysis of ligands of single MHC specificities (single allele (SA) ligands)(28–30) to computational motif deconvolution techniques(21,31,32) handling more complex multi-allele (MA) datasets. Within the latter category, the machine learning framework NNAlign_MA(33) has been demonstrated to efficiently deconvolute MA ligand data obtained from samples expressing multiple MHC alleles, enabling the construction of improved pan-specific predictors for antigen presentation for both the MHC class I and class II systems(33–35). NNAlign_MA achieves this by annotating the MA data during training in a semi-supervised manner based on MHC co-occurrence, MHC exclusion, and pan-specific binding prediction(33). This deconvolution expands the potential training data beyond binding affinity (BA) peptides and SA ligands to include the more complex and numerous MA ligands.

EL data differs from BA data in the sense that it not only captures peptide-MHC binding but also signals related to antigen processing. Recent MHCII prediction models(20,21,35) have leveraged these kinds of data and improved the prediction of MHCII antigen presentation.

Although most peptidome studies have focused on human and murine models, the technique can be equally applied to other species. In the context of livestock, we have earlier published studies demonstrating the ability to use mass spectrometry data to generate highly accurate prediction algorithms for BoLA-I molecules(36) which have been integrated into the NetMHCpan-4.1 server(34). Currently, there is no equivalent algorithm that can be used to predict peptide binding to BoLA-II molecules.

In this study, we have used mass-spectrometry to generate peptide elution data for BoLA-DR molecules and use the derived data to provide the first characterization of binding motifs of bovine MHCII and to demonstrate the development of the first available *in silico* method for accurate analysis of BoLA-DR ligands for rational CD4 T cell epitope prediction.

## Materials and Methods

### Animal and cell samples

Brazilian Holstein-Friesian PBMC samples were obtained from frozen archived materials from animals within the herd at the University of Sao Paulo that had been included in previous experiments completed under approval from the Committee on the Ethics of Animals Research at the Nowavet Veterinary Clinical Studies CRO, Viçosa/MG, certificate numbers 56/2016 (approved on 03 August 2016) and 36/2017 (approved on 09 June 2017). PBMC used for the characterization of BoLA-DR presented peptides from ovalbumin were isolated from a Holstein-Friesian animal from the University of Edinburgh herd with sampling conducted under a license granted under the UK Animal (Scientific Procedures) Act 1986. The *Theileria annulata-* and *Theileria parva*-infected cell lines used in this study had been established and characterised as part of previous studies and were maintained using routine and well-established protocols(37). The optimisation and final protocol used to assess the capacity of PBMC and *Theileria annulata*-infected cell lines to take up ovalbumin and present peptides on BoLA-DR molecules are described in Supplementary Figure 1.

### PBMC isolation, RNA extraction and cDNA synthesis

Bovine PBMC were isolated by density gradient centrifugation using Ficoll Paque Plus (GE Healthcare Bio-Sciences, Amersham. UK) according to manufacturers’ instructions. RNA was extracted from PBMC using TRIzol (Thermo Scientific, Renfrew, UK) and cDNA synthesised using the GOscript Kit (Promega, Southampton, UK), both according to the manufacturers’ instructions.

### BoLA-DRB3 sequencing

For BoLA-DRB3 amplification, primers (For - CCAGGGAGATCCAACCACATTTCC; Rev - TCGCCGCTGCACAGTGAAACTCTC) incorporating Illumina adaptors and multiplex identifier tags were obtained from IDT (Leuven, Belgium). PCR was performed using Phusion High Fidelity PCR kit (New England Biolabs), and the reaction was carried out in a final volume of 40 μL containing 2 μL of cDNA, 5X Phusion HF Buffer, 0.8 U μL of Phusion DNA Polymerase, 3% DMSO, 0.4 mM of dNTP and 0.5 μM of each primer. The reaction was performed in a G-Storm Thermal Cycle System (G-Storm) programmed for one cycle at 98 °C for 30 s, followed by 30 cycles at 98 °C for 10 s, 61 °C for 30 s, and 72 °C for 45 s, with a final extension period at 72 °C for 10 min. 5 μl of PCR product from each sample were pooled together, run on a 1.5% agarose gel, and the band of the appropriate size was extracted and purified using the QIAquick PCR Purification Kit (Qiagen). A final purification using Agencourt AMPure XP Beads (Beckman Coulter) at a ratio of 1:1 beads to PCR product was conducted prior to quantification of the sample and submission to Edinburgh Genomics for sequencing on the Illumina MiSeq V.3 platform. Analysis of the data was conducted using a bespoke bioinformatics pipeline (Vasoya *et al.* in preparation).

### pBoLA-DR complexes purification

Cultured cells (1×10^9^) were washed twice with ice-cold PBS and then lysed in buffer (1% IGEPAL, 15mM TRIS pH 8.0, 300 mM NaCl and cOmplete protease inhibitor (Roche)) at a density of 2×10^8^ cells/mL for 1 min, diluted with PBS 1:1 and solubilized for 45 min at 4 °C. Lysates were cleared by two-step centrifugation at 500g for 15 min at 4 °C and then at 15,000g for 45 min at 4 °C. For initial samples pBoLA-DR complexes were directly captured from the cleared lysates using 5 mg anti-BoLA-DR antibody (ILA21), immobilized in 1 mL of protein A resin (Amintra, Expedeon, Cambridge, UK). For later samples, pBoLA-DR complexes were captured from cleared lysates that had been depleted of peptide-BoLA-I (pBoLA-I) complexes by prior immunoprecipitation with 5 mg anti-BoLA-I antibody (ILA88), immobilized in 1 mL protein A resin. Captured pBoLA-DR complexes were washed, and peptides eluted from BoLA-DR molecules using 10% acetic acid and the resulting proteins dried as described in(38).

### HPLC

The dried pBoLA-DRB3 complexes were resuspended in 150 μL of loading buffer (0.1% formic acid, 1% acetonitrile) and loaded onto a 4.6 × 50 mm ProSwiftTM RP-1S column (Thermo Scientific) for reverse-phase chromatography on an Ultimate 3000 HPLC system (Thermo Scientific). Elution was performed using a 0.5 mL/min flow rate over 5 min on a gradient of 2 to 35% buffer B (0.1% formic acid in acetonitrile) in buffer A (0.1% formic acid). Eluted fractions were collected from 1 to 8.5 min, for 30 s each. Protein detection was performed at 280 nm. Even and odd eluted fractions were pooled together, vacuum dried and stored at −80 °C until use.

### LC-MS/MS

Dried samples were resuspended in 20 μL of loading buffer and analyzed in an Ultimate 3000 nano UPLC system online coupled to an Orbitrap Fusion™ Lumos™ Tribrid™ Mass Spectrometer (Lumos) (Thermo Scientific) or Q Exactive™ HF Hybrid Quadrupole-Orbitrap™ Mass Spectrometer (HFX). Peptides were separated in a 75 μm × 50 cm PepMap C18 column using a 1 h linear gradient from 2 to 30% buffer B in buffer A at a flow rate of 250 nL/min (~600 bar). Peptides were introduced into the mass spectrometer using a nano Easy Spray source (Thermo Scientific) at 2000 V. Subsequent isolation and higher energy C-trap dissociation (HCD) was induced in the 20 most abundant ions per full MS scan with an accumulation time of 120 ms and an isolation width of 1.2 Da (Lumos), or 1.6 Da (HFX). All fragmented precursor ions were actively excluded from repeated selection for 30 s. The mass spectrometry proteomics data have been deposited to the ProteomeXchange Consortium via the PRIDE(39) partner repository with the data set identifier PXDXXX (this ID will be made available upon manuscript acceptance).

### Mass spectrometry data analysis

The sequence interpretations of mass spectrometry spectra were performed using a database containing all bovine UniProt entries combined with entry P01012 for chicken ovalbumin (total of 41610 entries) and 4084 entries for *Theileria parva* Muguga proteome (40). The spectral interpretation was performed using *de novo*-assisted database search with PEAKS 10 (Bioinformatics Solutions), in ‘no enzyme’ mode, with mass tolerances of 5 ppm for precursor ions and 0.03 Da for fragment ions. The data was further searched against 313 inbuild peptide modifications.

### Filtering of MS-identified peptides

Previous to all analyses, the lists of peptides identified were filtered to remove: 1) peptides presenting post-translational modifications; 2) peptides with a peptide-spectrum matching score −Log10(P) < 15; 3) any peptides derived from *T. parva* Muguga, including the ones identified in both bovine and *T. parva* Muguga entries; and 4) peptides that shared a 9-mer overlap with the CD4 T-cell epitope benchmark.

### Model Training

All ligand data were filtered to include only peptides containing 13-21 residues, to exclude any residual potentially co-eluted MHCI peptides. Negative peptides were added as described earlier(35) by sampling random natural peptides from the bovine proteome (described below). Models were trained in a 5-fold cross-validation manner with partitions constructed from 9-mer common-motif clustering, ensuring no overlap between test- and training-data. Three model architectures were used (20, 40, and 60 hidden neurons), each trained with ten random weight initialization, resulting in an ensemble of 150 networks. Models were evaluated in a percentile rank fashion, meaning that prediction scores are normalized against a distribution of prediction scores from random natural peptides. Rank scores are more interpretable than raw prediction scores and allow for fairer comparison across alleles.

Two models were trained in this project, both using the NNAlign_MA machine learning framework(33). The first model (BoLA) was trained on the novel BoLA SA and MA EL data combined with the BA data from NetMHCIIpan-4.0 with an added set of BoLA BA data (roughly 250 measurements for each BoLA-DR molecules incorporating the three different BoLA-DRB3 alleles - generated *in house).* For the second model (All Data), the BoLA EL data were combined with all the EL data from the NetMHCIIpan-4.0 data set (human and murine EL data) and the same BA data as the BoLA model. The BoLA and All Data models share partitions.

Explicit encoding of ligand context was leveraged to capture antigen processing signatures, as previously described (20). Briefly, in context encoding 12 residues of the ligand and antigen are fed as input to the model, 6 are from the N-terminal region of the ligand (3 residues upstream of the ligand in the antigen and 3 N-terminal Peptide Flanking Regions (PFRs)), and 6 are from the C-terminal region (3 C-terminal PFRs and 3 downstream of the ligand).

Peptide lists resulting from BoLA-DR eluted ligand data are by nature only positive examples of ligands that interact with MHCII (excepting co-eluting peptide noise from assay). To train a peptide-MHCII interaction model, the training data must include examples of non-interacting peptides sampled from the same background as positive data. To achieve this, peptides (and their context, see above) were randomly sampled from the bovine proteome. Random negative peptides were made to follow a uniform length distribution of 13-21 residues, sampling for each length five times the number of peptides in the most commonly observed ligand length for a dataset. Negatives were sampled independently for each bovine dataset with a uniform length distribution so the model can learn the length distribution of ligands(27,41).

## Results

### Analysis of the BoLA-DRB3 repertoire in an experimental cohort of Brazilian Holstein-Friesians

The IPD-MHC database includes over 300 BoLA-DRB3 alleles, of which only a small subset could be included in this study. To identify the alleles that would be most relevant to ongoing experiments, a novel high-throughput MiSeq BoLA-DRB3 sequencing approach (Vasoya *et al.,* in preparation) was used to examine the frequency of DRB3 alleles in a representative cohort of 30 Holstein-Friesian animals from the experimental herd at the University of São Paulo, Brazil. A total of 22 DRB3 alleles were identified, including a novel allele that had not been previously described (nDRB3.1). Typical of MHC allele distribution in most cattle populations, there was a small number of dominant alleles, DRB3*15:01, DRB3*01:01, DRB3*11:01, DRB3*14:01:01, and DRB3*12:01, which were present at a frequency of ≥5%, whilst the remaining 17 alleles were present at lower frequencies (Figure 1).

**Figure 1.**
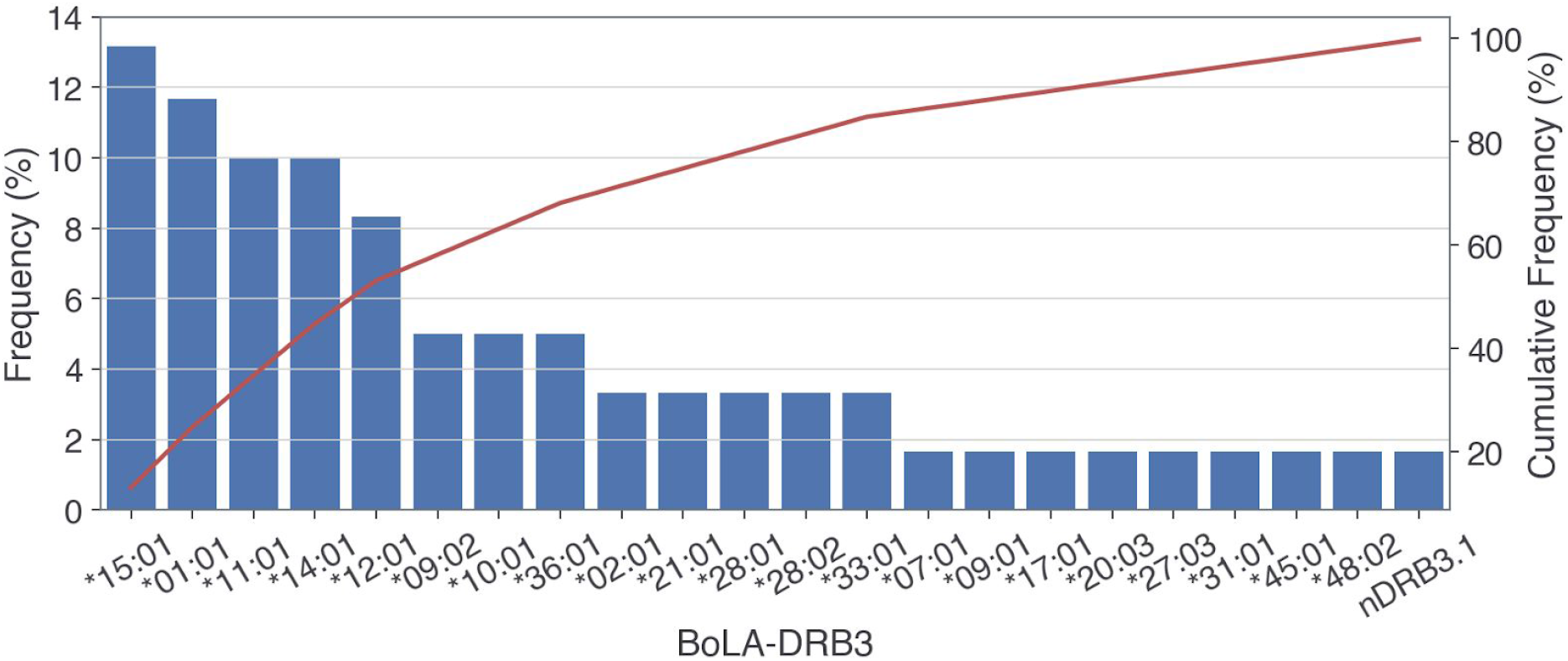
Frequencies of BoLA-DRB3 alleles detected by a MiSeq genotyping approach in a subset of the experimental Holstein-Friesian cattle herd at the University of Sao Paulo (n=30). The frequency data is shown as a Pareto plot with the frequency of individual alleles displayed on the left vertical axis and the cumulative frequencies of the DRB3 alleles shown on the right vertical axis. Allele nDRB3.1 was a novel sequence.

### Generation and analysis of MS data for BoLA-DR eluted peptides

Initial experiments to establish a BoLA-DR elution technique used O11 and 2229 *Theileria annulata* (TA) cell lines which had previously been confirmed to be homozygous for DRB3*10:01 and DRB3*11:01, respectively (Table 1). The length distribution of the peptides obtained from the 2229TA and both replicates (n1 and n2) of O11TA cell lines was bi-modal. One peak, centred around 14-15mers was the size anticipated for MHCII ligands; the second peak, centred around 8-10mer peptides, was more consistent with the length distribution of MHCI ligands (Figure 2A), and it was speculated that this represented a substantial level of co-purification of BoLA-I molecules during BoLA-DR immunoprecipitation. To investigate this, NetMHCpan-4.1(34) was used to predict the binding potential of all 8-13-mer peptides in each of the MS data sets for each of the BoLA-I molecules expressed in the given cell line (Table 1). The sequence logos of these peptide sets (Supplementary Figure 2) showed remarkable similarity to the motifs previously described for the BoLA-I alleles in these haplotypes(34) and between 56.8-70.9% of the 8-13-mer peptides in each sample were predicted to be BoLA-I binders (defined using a binding threshold of 5% rank). This corroborated the hypothesis that the majority of these peptides originated from co-precipitated BoLA-I ligands and their removal resulted in a substantial diminution of the 8-10mer peak (Figure 2B).

**Figure 2.**
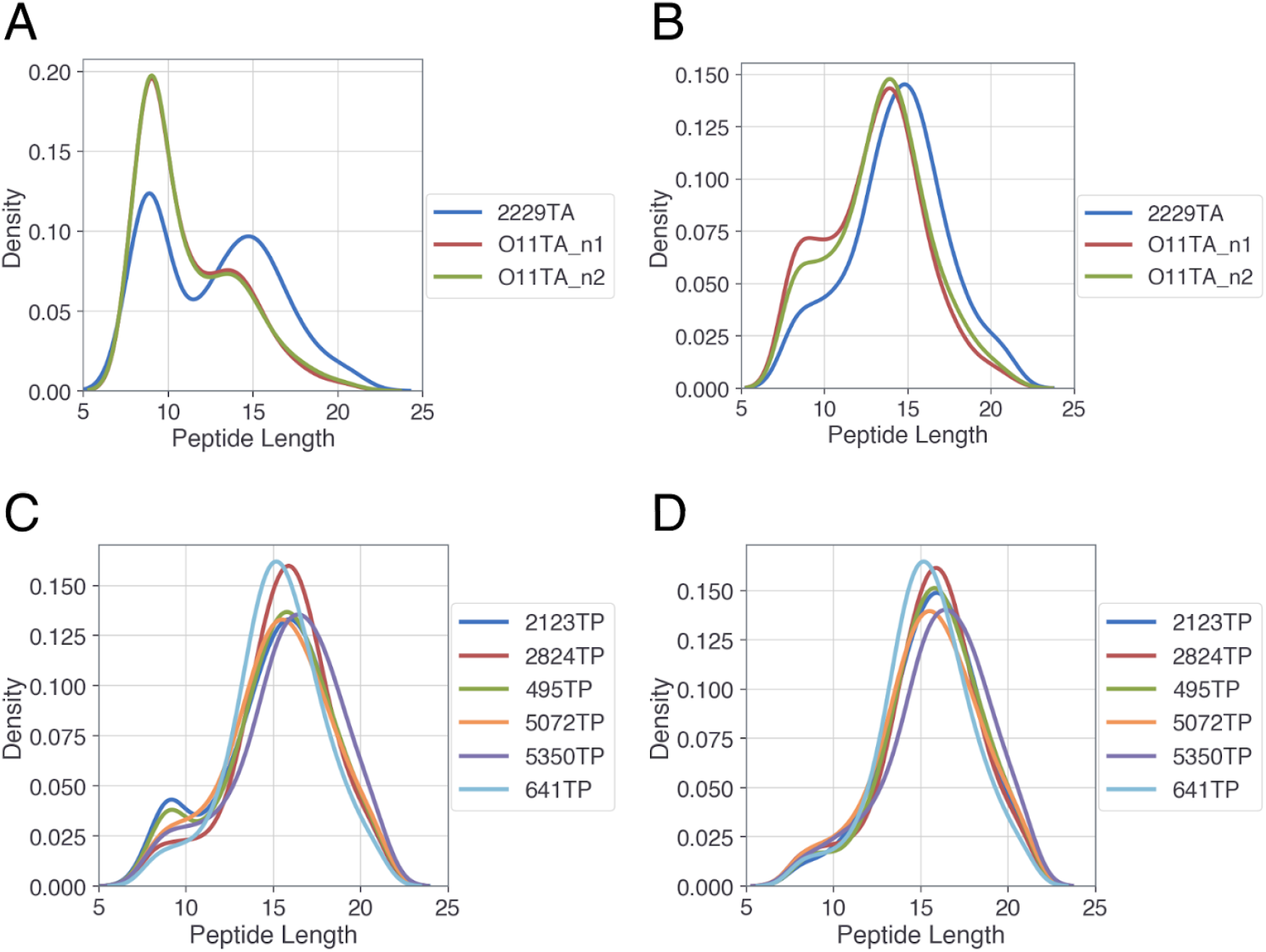
Length distribution of BoLA-DR eluted peptides. Kernel density estimates comparing length distributions of BoLA-DR eluted peptides using different strategies for removal of BoLA-I eluted contaminants: (A) Direct pBoLA-DR elution; (B) Direct pBoLA-DR elution with subsequent removal of BoLA-I binders as predicted by NetMHCpan-4.1; (C) Initial immunoprecipitation to deplete pBoLA-I complexes. (D) Same as for panel (C) but with subsequent removal of BoLA-I binders as predicted by NetMHCpan-4.1. Due to failed pBoLA-I depletion sample 2229TP is not represented in this figure.

**Table 1.**
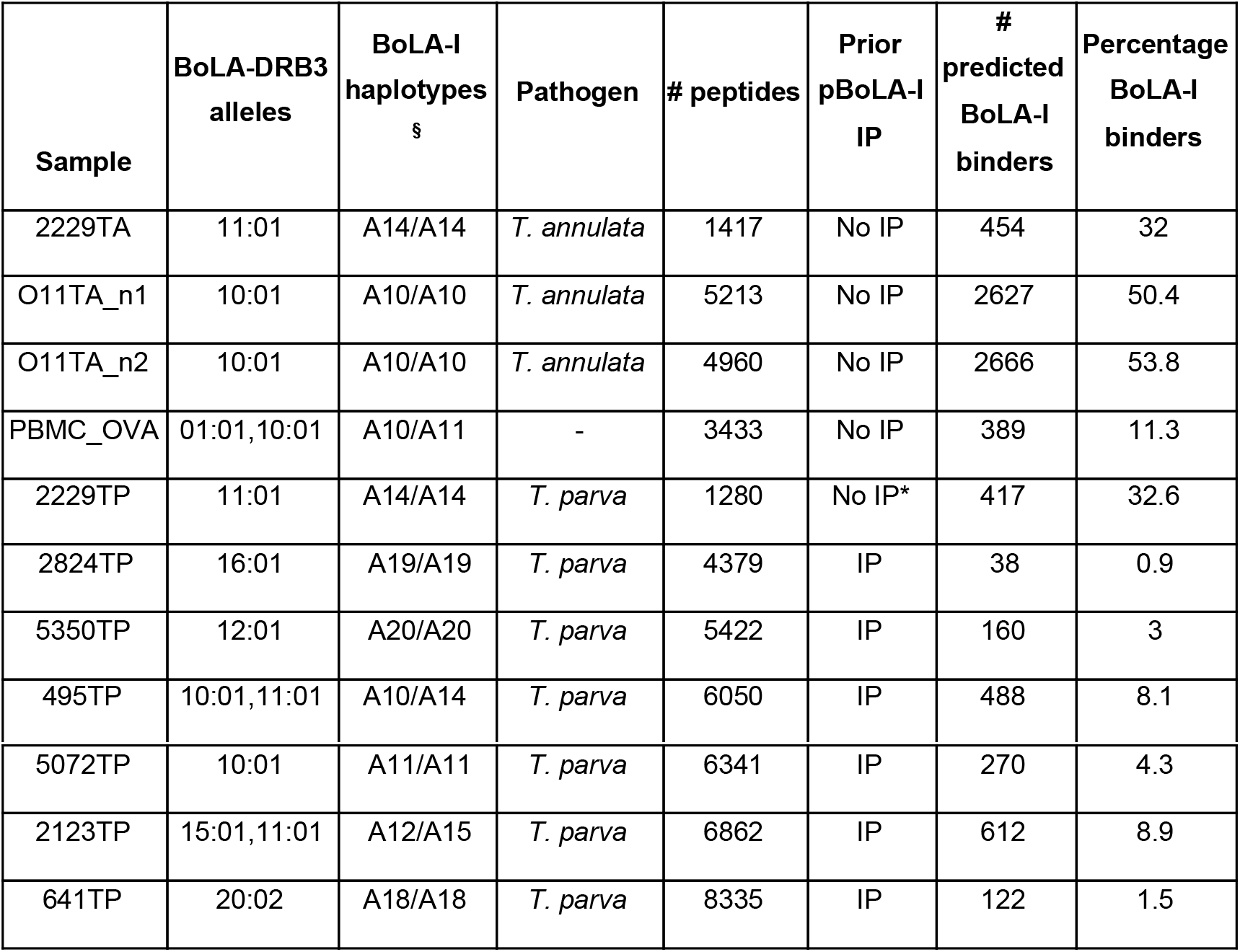
Overview of MS BoLA-DRB3 ligand elution datasets. For each sample, information regarding BoLA-DRB3 alleles, BoLA-I haplotypes, infecting pathogen, the number of ligands identified in each sample, use of prior pBoLA-I immunoprecipitation (IP) and number and percentage of predicted BoLA-I binders are shown. The 2229TA and O11TA samples were generated without prior pBoLA-I immunoprecipitation (see text). Samples PBMC_OVA and O11TA_n2 were OVA-loaded. *Preliminary pBoLA-I IP depletion failed on this sample. §: from https://www.ebi.ac.uk/ipd/mhc/group/BoLA/haplotype/

To address the observed co-enrichment of pBoLA-I in pBoLA-DR immunoprecipitations, it was decided to apply a sequential immunoprecipitation protocol, starting with pBoLA-I complex depletion using an anti-BoLA-I monoclonal antibody (IL-A88), followed by pBoLA-DR precipitation. This two-step protocol was applied to samples from a series of seven *T. parva*-infected cell lines (Table 1) which expressed a range of DRB3 alleles present in our experimental cohort (*11:01, *10:01, *1501, *1201) or which were of interest because of ongoing *T. parva* CD4 T cell epitope identification studies that included these alleles (*16:01 and *20:01). The total numbers of peptides identified in these samples ranged between 1280 and 8335 (Table 1), and the distribution of the peptide lengths is shown in Figure 2C. The results in this figure show a substantially lower representation of 8-10mer peptides, indicating successful reduction but not complete depletion of BoLA-I eluted peptides (Figure 2C). Analysis of the binding potential of the peptides in the 8-10mer peak confirmed that the majority were, in fact, still BoLA-I binders (Table 1 and Figure 2D); indicating that although the preliminary BoLA-I depletion had a profound effect on reducing peptides from co-eluted pBoLA-I, it did not eliminate them completely. Removal of predicted MHCI binders from the datasets (ranging in frequency from 0.9-8.9%, Table 1) effectively abolished the 8-10mer peak (Figure 2D), establishing that i) combined BoLA-I depletion by prior immunoprecipitation and bioinformatic removal of predicted MHCI-binders provided the optimal results and ii) consistent with other MHCII molecules, BoLA-DRB3 molecules have a preference for binding peptides of length 13-21 amino acids (after the combined filtering, 80.7% of the peptides fall in this length range).

### Motif deconvolution and prediction model generation from MS data sets of BoLA-DR eluted ligands

Using the MS BoLA-DR EL data sets, alternative models for BoLA-DRB3 motif deconvolution were assessed and a prediction model for BoLA-DRB3 ligands was developed. Details for the model training and model parameters are described in the materials and methods. In short, bovine ligand data was filtered only to include peptides of 13-21 residues and were used as positive data points, with negative data points added as previously described(35). Two models were trained: a ‘BoLA’ model using the novel BoLA-DR elution data combined with the BA (binding affinity) data from NetMHCIIpan-4.0 and a set of BA data covering three different BoLA-DRB3 alleles; and an ‘All Data’ model, which includes the BA and EL data of the BoLA model with added murine and human EL data from the NetMHCIIpan-4.0 data set. Both models were trained with and without assessing the ‘context’ of the peptide within the parent protein (MAC- and MA-models, respectively). Here, ligand context refers to including residues near the ligand termini, inside and outside the ligand, to capture signals of antigen processing. Further details on data partitioning, model training and context definition are provided in materials and methods.

The results of the cross-validation evaluation measured in terms of the AUC are shown in Figure 3 and show clear differences in the performance of the models used. Firstly, for both the ‘BoLA’ and the ‘All Data’ models, every cell line data set displayed a higher AUC for the MAC-model than the MA-Model (p-value: 0.00097 in a binomial test counting number of cell lines with higher AUC for MAC-models versus MA-models). This agrees with earlier studies for the human and mouse MHCII system(20,35,42), showing the value of incorporating encoding context into the prediction models. Secondly, the ‘BoLA’ MAC-model has significantly higher median AUC compared to the ‘All Data’ MAC-Model (p-value: 0.00195 in a binomial test counting cell lines where ‘BoLA’ MAC-model has higher AUC compared to ‘All Data’ MAC-model, excluding ties), indicating that inclusion of the human and murine training data had no benefit in the generation of a model for BoLA-DR binding prediction. This comparative evaluation clearly demonstrated the ‘BoLA-MAC’ model exhibited the best performance and so was selected for subsequent use.

**Figure 3.**
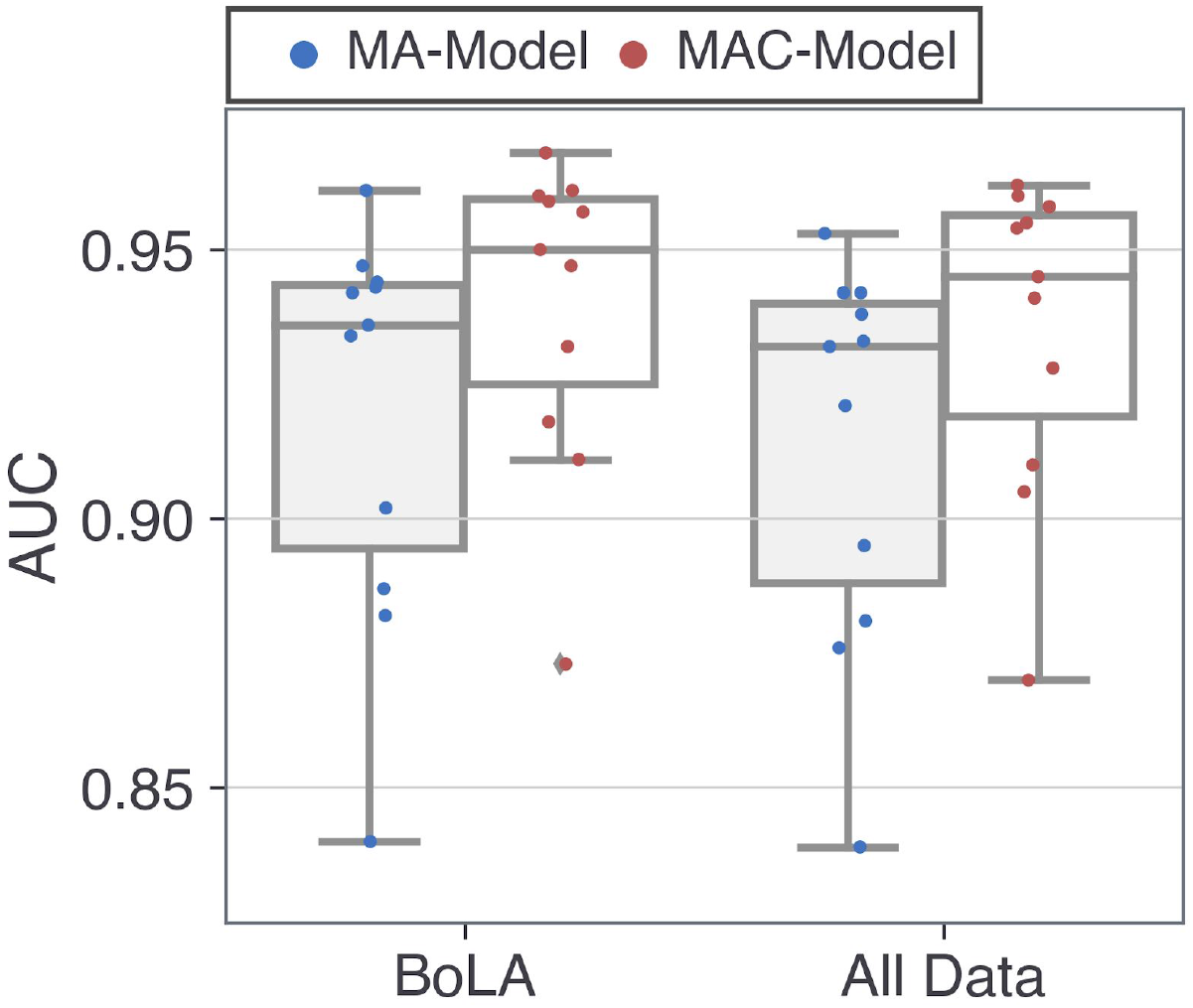
Cross-Validation evaluation of bovine EL data. Models were evaluated on the BoLA-DR ligand data in a cross-validation manner. The boxplot shows the AUC per cell line sample for the BoLA and All Data models with and without context encoding (MAC-Model and MA-Model, respectively). Each point in the figure represents data from a single sample. Of note, the outlier sample with a cross-validated AUC performance below 0.90 for the BoLA-MAC model was 2229TA; this sample had 27% ligands assigned as contaminants causing the decrease in the observed AUC (Supplementary Figure 3).

Examples of BoLA-DRB3 allele motif deconvolution from EL data-sets as performed by the BoLA-MAC model are shown in Figure 4. The motif deconvolution results for each sample included in this study are displayed in Supplementary Figure 3, and the motifs for each of the seven BoLA-DRB3 alleles covered by the EL data (combining the data from all samples) are shown in Supplementary Figure 4. As can be seen in Figure 4, the deconvolution results in well-defined motifs, with the anticipated preference for residues at positions 1, 4, 6 and 9 of the binding core and limited exclusion of non-conforming peptides (average of 8.6% of ligands assigned as contaminants in samples included in Figure 4). The data presented here also shows the ability of the deconvolution to discriminate the motifs of both BoLA-DRB3 alleles in heterozygous samples (495TP and 2123TP) as well as the consistency in the motifs for the same BoLA-DRB3 molecule obtained from different EL data-sets (e.g. BoLA-DRB3*10:01 in 495TP and 5072TP). These observations are consistent across all of the samples included in this study, with non-conforming (trash) peptides constituting only ~12.5%, a high average Pearson correlation between motifs for the same BoLA-DRB3 molecule (0.92 for BoLA-DRB3*10:01 and 0.908 for BoLA-DRB3*11:01, Supplementary Figure 5), and a very high specificity being demonstrated for individual motifs (PPV values in the range 0.751-0.868, across the different deconvolutions, Supplementary Table 1). As such, the data confirms that the BoLA-MAC model permitted the generation of high resolution and reproducible BoLA-DRB3 binding motifs from EL data. This model, renamed as NetBoLAIIpan, has been made publicly available at www.cbs.dtu.dk/services/NetBoLAIIpan.

**Figure 4.**
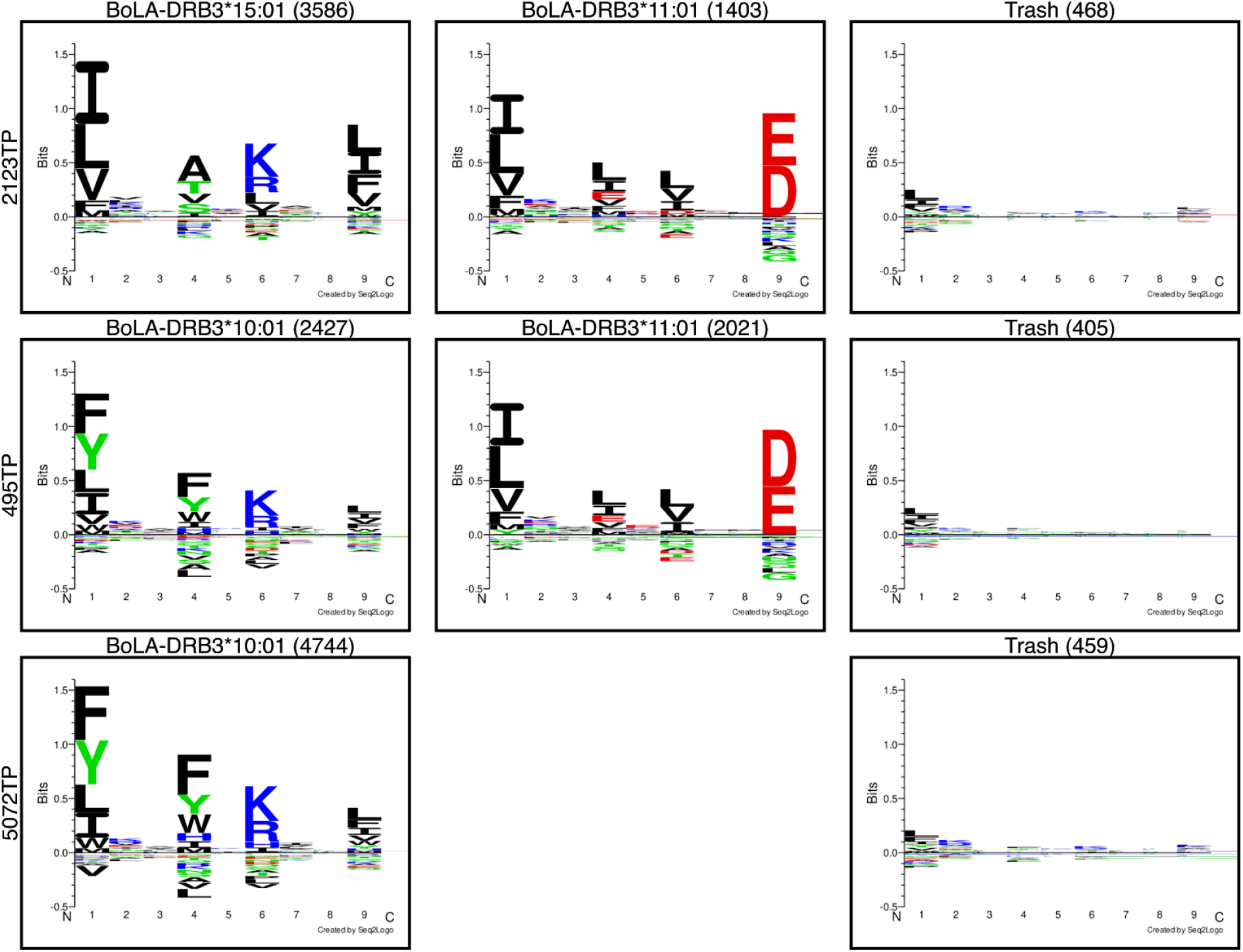
Examples of deconvoluted motifs derived from EL BoLA-DR datasets. From each cell line defined as being heterozygous for DRB3, two peptide-binding motifs were derived. Where cell lines express the same DRB3 allele, consistent motifs were identified (e.g., both 2123TP and 495TP express DRB3*11:01 and show a similar peptide-binding motif). Motifs were generated from ligands with a rank score of < 20 for the context-model. Ligands with a predicted rank >20 are assigned to the Trash cluster. Logos show alignments of predicted peptide binding cores where numbers in parenthesis represent the number of binding cores.

### NetBoLAIIpan can be used to predict BoLA-DRB3 presented peptides derived from exogenous proteins

To extend our studies on the utility of the NetBoLAIIpan method developed above, the model’s ability to predict which peptides would be presented by BoLA-DR molecules from an exogenous protein was examined. Here, both PBMC (BoLA-DRB3*01:01 and *11:01) and the O11TA_n2 cell line (BoLA-DRB3*10:01) described above were pulsed with soluble ovalbumin (OVA, see materials and methods and Supplementary Figure 1 for details) before performing pBoLA-DR elution. Only one OVA-derived peptide (“SSANLSGISSAESLK”) was identified in the O11TA sample, which demonstrated very poor predicted binding to BoLA-DRB3*10:01 with a predicted percentile rank value of 29.2%, strongly suggesting it was not a genuine BoLA-DR binding peptide. In contrast, seven OVA-derived peptides were identified in the PBMC sample. Mapping the seven peptides onto the OVA protein sequence (Figure 5 - Inserted panel) shows that all the peptides clustered around the 9-mer core “INKVVRFDK”, located at OVA_54-62_, with a common motif IxxVxRxxK - matching the motif described in Supplementary Figure 4 for BoLA-DRB3*01:01. Also of interest is that six out of the seven ligands observed had proline in the C-2 position, which is a common feature in context motifs(20). The NetBoLAIIpan model was applied to predict potential DRB3*01:01 and DRB3*10:01 ligands in the OVA protein sequence. To achieve this, the OVA protein was *in silico* digested into overlapping 13-21-mer peptides, and binding to DRB3*01:01 and DRB3*10:01 was predicted for each peptide with predicted ligands identified using a 1% rank score threshold; this resulted in the identification of 48 predicted ligands. The MS identified and *in silico* predicted ligands were then stacked onto the OVA protein sequence, and a profile was calculated showing the relative number of measured and predicted ligands mapped to each amino acid position within the protein. The MS identified and *in silico* predicted ligand profiles demonstrated a striking concordance, with the MS identified peptides overlapping with the dominant peak of *in silico* predicted peptides (38 overlapping peptides located at positions 45-71) (Figure 5) (similar data were obtained using rank threshold values in the range 0.5-2.0%, results not shown), indicating that NetBoLAIIpan can accurately predict ligands derived from defined proteins that are experimentally shown by MS to be presented by BoLA-DR.

**Figure 5.**
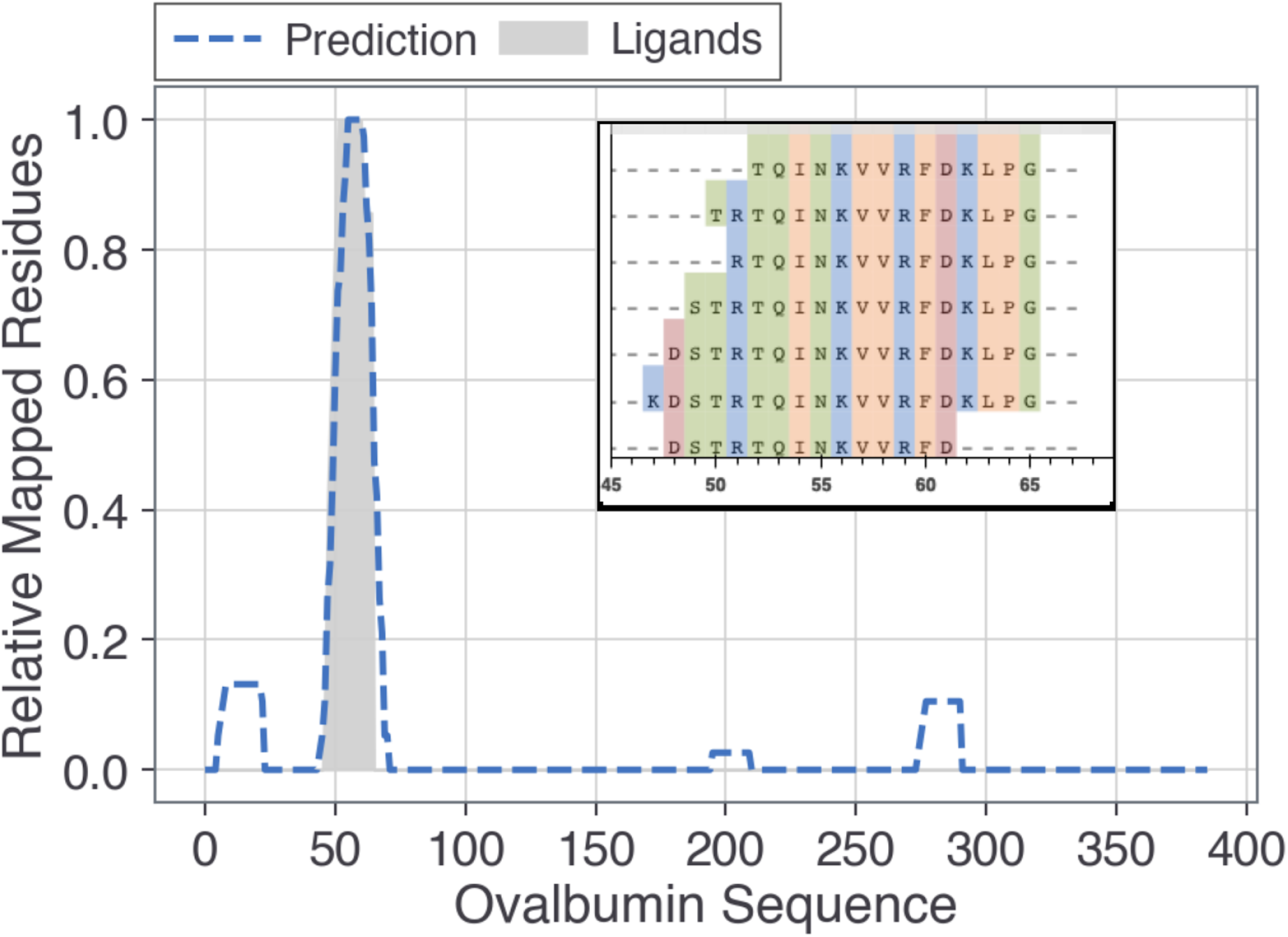
Profiles of predicted and measured OVA ligands in the PBMC cell line. (Main Figure) The gray shaded area shows the relative number of measured EL ligands in the PBMC sample overlapping each position in the OVA sequence. The dotted line represents the mapping of 13-21-mers from the OVA sequence predicted with a rank score < 1% for the BoLA-DRs expressed in the PBMC sample; the peaks at positions 6-23, 45-71, 196-210 and 275-291 represent 5, 38, 1 and 4 predicted BoLA-DR binding peptides, each with median predicted rank scores of 0.64, 0.45, 0.82, and 0.56, respectively. (**Inserted panel)** Mapping of the seven OVA peptides measured in the PBMC cell line. All but one of the peptides shared a binding core “INKVVRFDK” in positions 54-62 of the OVA sequence.

### Validation of the BoLA model for BoLA-DRB3 presented CD4 T cell epitope prediction

Next, the performance of NetBoLAIIpan was validated using a set of 25 experimentally validated BoLA-DR restricted *T. parva* CD4 T cell epitopes (Morrison et al., manuscript in preparation, refer to Supplementary Table 2). Here, the NetMHCIIpan-4.0 was included as a reference model to test the extent to which peptide presentation rules learned from human and murine data extrapolate to bovine epitopes. Each epitope source protein was *in silico* digested into peptide strings matching the length of the epitopes, and each peptide was then assigned the lowest predicted rank score from the set of 13-19-mers whose binding core overlapped with the peptide string. Next, the epitope’s F-rank value was calculated as the percentage of peptides with a greater prediction score than the epitope. Hence, a perfect prediction has an F-rank value of 0, and a random prediction presents a value of 50. Comparison of F-rank values obtained by the different models for the set of *T. parva* epitopes (Figure 6), shows that the NetBoLAIIpan models with or without context achieved equivalent prediction performance both achieving a median F-rank value of 0.697% and median prediction percentile rank score for the epitopes of 0.2. In practical terms, these results translate into 12 out of 25 epitopes being ranked as the top predicted peptide within the given source protein. Both NetBoLAIIpan models achieved significantly better F-ranks compared to NetMHCIIpan-4.0 (p-values: <0.001 comparing the two NetBoLAIIpan models to NetMHCIIpan-4.0). The large difference in the performance of the NetMHCIIpan-4.0 and NetBoLAIIpan models clearly demonstrates the power of combining BoLA-DR EL data and advanced immunoinformatics to generate novel tools for characterizing antigen presentation epitope identification in the BoLA-DR system.

**Figure 6.**
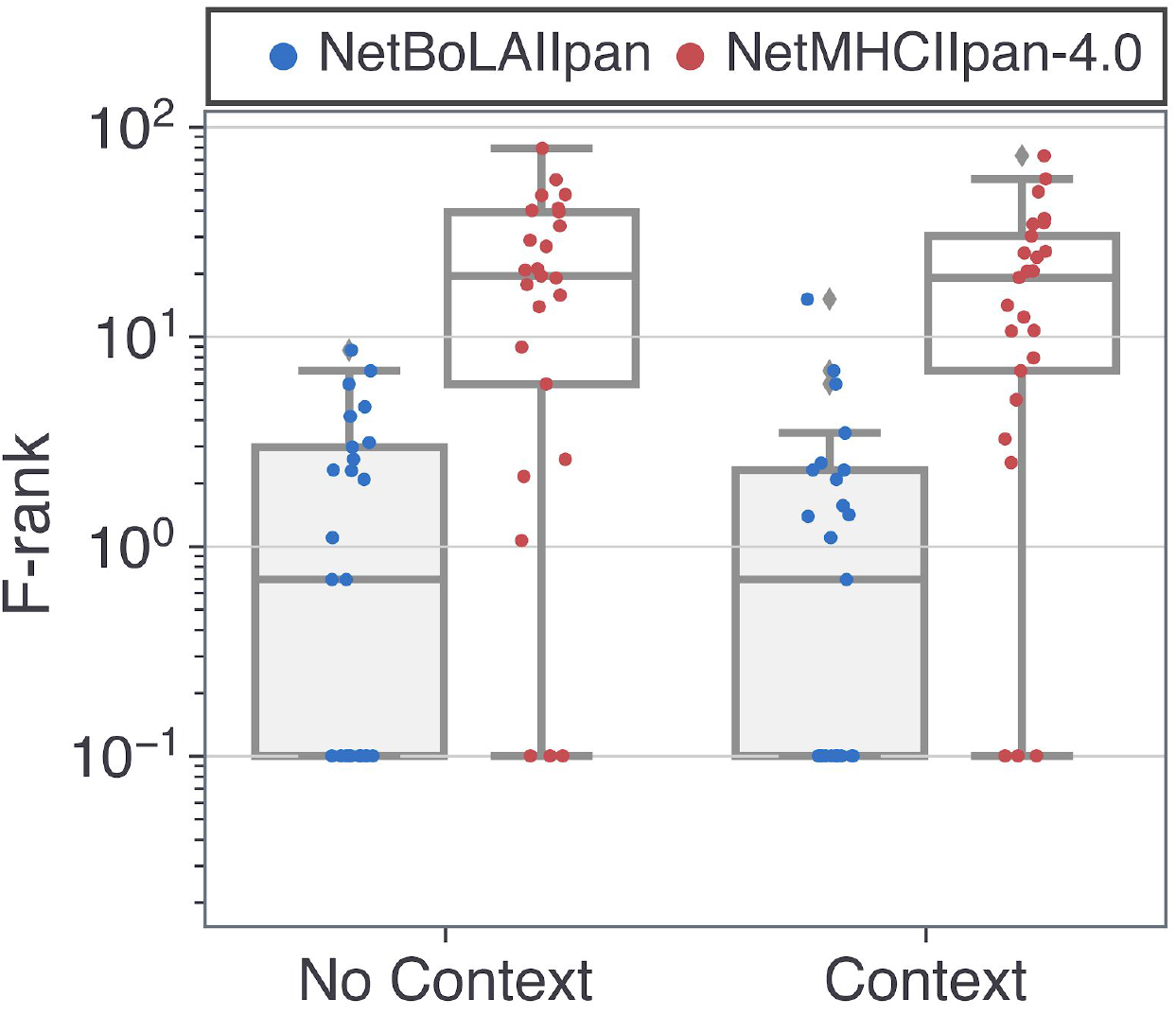
Comparison of different BoLA-DR prediction models using validated CD4 T cell epitopes. Distribution of percentage F-rank performance values for defined BoLA-DR presented *T. parva* epitopes using the NetBoLAIIpan and NetMHCIIpan-4.0 models with (Context) and without context (No Context). Prediction scores were assigned to each overlapping epitope length-matched peptide in the epitope source protein as described in the text. The y-axis is shown in log-scale and F-rank values below 0.1 are presented as 0.1005 to avoid non-defined values.

## Discussion

A pre-requisite for the development of next-generation subunit vaccines is the identification of antigens containing epitopes that can be recognised by B cells, CD8 T cells and CD4 T cells, as appropriate for the immune response required. Several bioinformatic tools that enable the prediction of CD4 T cell epitopes in humans have been developed and the recent integration of large-scale MHC-eluted peptide data have led to a dramatic improvement in their performance(21,30,35). In contrast, there is a lack of equivalent bioinformatics tools designed specifically for bovine MHCII molecules, and since the currently available tools have not incorporated bovine MHCII EL data during their development, they perform with limited accuracy when applied to bovine data (as demonstrated in this study - Figure 6). In previous studies, we have shown how the use of high-quality EL mass spectrometry data combined with advanced immunoinformatics and machine-learning techniques can further our understanding of the rules underlying MHC antigen processing and presentation, allowing the development of improved prediction methods for MHC ligands and T cell epitopes(33-35). Here, we have extended this work to cover, for the first time, BoLA-DR molecules.

Results from our initial experiments indicated that the peptides isolated following pBoLA-DR immunoprecipitation were heavily contaminated with co-eluted pBoLA-I-presented peptides. This phenomenon has been reported previously in other studies using equivalent protocols for immunoprecipitation of MHCII molecules from human cell lines and has been hypothesised to reflect that the protocol for lysing the cells results in the immunoprecipitation of membrane fractions, which contain both MHCI and MHCII molecules(43,44). In this study neither prior depletion of pBoLA-I (by immunoprecipitation) nor bioinformatic prediction and removal of BoLA-I contaminant ligands were completely effective in eliminating the BoLA-I-binding contamination when applied alone - both left a remnant peak of 8-10-mer peptides. However, the combined use of these two approaches was successful in removing the 8-10-mer peptide peak, resulting in 13-21-mer dominated profiles characteristic of MHCII presented peptides. On this basis we would propose that future studies for BoLA-II immuno-peptidomics should routinely make use of both preliminary depletion of pBoLA-I complexes by use of an initial pBoLA-I immunoprecipitation step (consistent with recently developed approaches for human MHCII immuno-peptidomic studies(21,45)), and *in silico* immunoinformatic BoLA-I peptide-binding depletion using currently available prediction methods(34,36) (or if working with cell lines expressing alternative BoLA-I haplotypes by generating BoLA-I peptide-binding motifs by subjecting the product of the preliminary pBoLA-I immunoprecipitation to elution, mass-spectrometric analysis and subsequent motif deconvolution).

In this study, we compared two models for developing the BoLA-DR prediction algorithm. The first of these was trained using EL data only from BoLA-DR, whilst the second was trained on the same data augmented by an exhaustive human (HLA) and murine (H-2) MHCII-eluted peptide dataset (both models also incorporated human, murine and a small amount of bovine BA data). A cross-validation evaluation demonstrated that the former model had superior performance, suggesting that integration of cross-species EL datasets was not beneficial to the accuracy of the results generated by this model. However, this evaluation was restricted to the limited set of BoLA-DRB3 alleles covered by the EL data generated in the current study, and it remains to be seen whether a model integrating cross-species EL data would allow improved prediction when extrapolated to data generated from samples expressing other BoLA-DRB3 alleles. As over 300 BoLA-DRB3 alleles have been described at present, further evaluation of how best to incorporate inter- and intra-species data to improve the algorithm’s performance is warranted as it will not be feasible for BoLA-DR EL data to be generated for more than a subset of these alleles. The seven BoLA-DRB3 alleles included in this study were selected predominantly based on their frequency in the experimental herd of Holstein-Friesian cattle at the University of São Paulo (USP) (in combination with the availability of DRB3-genotyped TA/TP cell lines and validated BoLA-DRB3 presented epitope data). The cumulative total frequency of these seven alleles in the samples of animals from the USP herd was ~48% and retrospective analysis of the University of Edinburgh herd shows that these alleles have an even higher representation (~67.9%). This is broadly in line with the frequencies observed in Holstein-Friesian herds across South America and other parts of the world (51.2-73%)(18). Analysis of the BoLA-DRB3 molecules in Holstein-Friesian animals is attractive for several reasons: i) due to the high levels of inbreeding, characterisation of a small number of DRB3 alleles will allow comprehensive coverage of the breed (e.g. inclusion of another five DRB3 alleles would give 77-98% coverage of Holstein-Friesian populations(18) and ii) as high-value dairy animals there is great interest in introducing Holstein-Friesians into low-income countries (frequently tropical) as part of the process of increasing agricultural productivity and food security; a major limitation to this process is the Holstein-Friesian susceptibility to many of the pathogens prevalent in regions of the world. Consequently, there is a particular interest in finding interventions, such as vaccination, that can be used to protect Holstein-Friesian animals in tropical environments.

A critical and general issue for rational vaccine development is the identification of relevant antigens. Approaches dependent on conventional antigen-screening techniques have limitations, especially when applied to complex pathogens (e.g. eukaryotic pathogens), where the size of the proteomes makes a comprehensive analysis of the full potential antigen repertoire prohibitively expensive and laborious. For such pathogens, bioinformatic tools that can help rationalise antigen screening assays and/or selection are of particular value and have a significant potential for accelerating vaccine development. A potential approach would be to use bioinformatics tools to predict which peptides from a candidate antigen would be present by BoLA molecules when delivered as a vaccine. To directly evaluate this, we examined NetBoLAIIpan’s ability to correctly identify the peptides from ovalbumin that had been pre-loaded onto cell’s then subjected to MHC-elution analysis. A comparison of the set of eluted peptides from a PBMC sample and the *in silico* predicted BoLA-DRB3 binding peptides demonstrated an exceptionally high level of concordance. This suggests that the ability of NetBoLAIIpan to accurately model the peptides derived from an exogenously administered protein could be exploited to provide an efficient and inexpensive *in silico* preliminary evaluation of the potential immunogenicity of candidate antigens and so contribute to the rational selection of antigens(46) prior to undertaking expensive and laborious *in vivo/in vitro* experiments. In particular, such an analysis could be used to assess the MHC coverage of individual antigens, and thus inform the construction of optimal vaccine designs. An example of how such *in silico* analysis could be employed is given in Supplementary File 6.

During the development of the prediction model, it was clear that the integration of signals relating to antigen-processing was beneficial. That is, the inclusion of information regarding the ‘context’ of the peptides (i.e. both the amino acid residues in the protein flanking the peptides and the amino acids at the termini of the peptide) significantly improved the power of the models for predicting ligands. The NetBoLAIIpan model exhibited an unprecedented high performance when evaluated using a set of validated BoLA-DRB3 presented epitopes from *T. parva,* achieving a median F-rank score of 0.697% (corresponding to 12 out of 25 of the defined epitopes being the highest predicted peptides within the source protein). This performance was significantly higher than the 19.23% achieved by the previously available NetMHCIIpan model which had not been trained on the BoLA-DRB3 elution peptide data, demonstrating the utility of generating and incorporating these data sets. In line with earlier work, context did not impart the same benefit in the task of ranking CD4 epitopes as was found for ligand data. Here, the context model was found to perform equivalent to the non-context model. These results align with earlier work using the mouse and human MHC class II systems(20,35,42). Interestingly, however further improvements in epitope prediction could be obtained by ranking antigen peptides based on the number of binders within overlapping 13-19-mers. This method of assigning epitope ranks is based on the intuitive assumption that protein regions with multiple predicted binders have a greater chance of being presented by BoLA-DRB3 molecules. Using this approach, the median F-rank score was 0.362%, suggesting a non-trivial improvement in the prediction. However, further benchmarking on larger epitope sets to systematically evaluate the comparative performance of this methodology is needed before the recommendation that it is routinely adopted can be made.

In conclusion, this study has proven the high value and important synergistic effect of combining peptide-MHC elution MS data and advanced immunoinformatics to characterize antigen presentation and perform ligand/epitope identification in the BoLA-DR system.

## Acknowledgments

This work was supported in part by funds from the National Institute of Allergy and Infectious Diseases, National Institutes of Health, Department of Health and Human Services, under Contract No. HHSN272201200010C and from the Fundação de Amparo à Pesquisa do Estado de São Paulo - FAPESP (2015/09683-9) to BRF, the Bill & Melinda Gates Foundation and with UK aid from the UK Foreign, Commonwealth and Development Office (Grant Agreement OPP1127286) under the auspices of the Centre for Tropical Livestock Genetics and Health (CTLGH), established jointly by the University of Edinburgh, SRUC (Scotland’s Rural College), and the International Livestock Research Institute (the findings and conclusions contained within are those of the authors and do not necessarily reflect positions or policies of the Bill & Melinda Gates Foundation nor the UK Government) and the BBSRC through the ISP award made to The Roslin Institute (BBS/E/D/20002174). AF was supported by FAPESP scholarships 2014/11010-9, 2017/21401-4 and 2018/23579-8.

