## Supplementary Materials for "Integral use of immunopeptidomics and immunoinformatics for the characterization of antigen presentation and rational identification of BoLA-DR-presented peptides and epitopes"

**Supplementary Figure 1 - Experimental optimization for exogenous protein presentation assay**

***Optimisation of the conditions for loading ovalbumin (OVA) as a model protein onto BoLA-DR molecules:*** in preliminary small scale studies fluorescent ovalbumin (fOVA) was used to optimise the conditions for ovalbumin loading into *T. annulata*-infected cells and PBMC. fOVA at a range of concentrations (125-750µg/ml) was incubated with *T. annulata*-infected cells at a concentration of 10^7 cells/ml *in vitro* for either 3 h or 6 h at either 4°C or 37 °C. All incubations were completed in standard culture media composed of RPMI 1640 supplemented with 10% heat-inactivated fetal bovine serum, 2 mM L-glutamine, 100 U/ml penicillin/streptomycin (Thermo Scientific), and 50 µM 2-mercaptoethanol (Sigma Aldrich). Previous analysis had shown that >99% of the *T. annulata*-infected cells expressed BoLA-DR on the cell surface (as measured using the ILA21 antibody). After the period of incubation the cells were washed twice in FACS buffer (PBS + 0.5% BSA, 2 mM EDTA, 0.05% sodium azide), and the uptake of fOVA measured using an LSR Fortessa X20 Cell Analyzer (Becton Dickinson). Incubation of the *T. annulata*-infected cells for 6 h with ≥ 500µg/ml of fluorescein-conjugated ovalbumin (fOVA) resulted in ~99% of *T.annulata* cells having internalised fOVA (Supplementary Figure 1a). fOVA internalisation was much reduced when the cells were incubated at 4°C indicating the uptake of fOVA was a metabolically active process. PBMCs were cultured in the same conditions for 6 h with 500µg/ml fOVA and then stained with Zombie Yellow viability dye (1:1000, Biolegend, London, UK), followed by ILA21 (murine IgG2a - 1µg/mL) and goat anti-mouse IgG2a Alexa Fluor 647 (1:2000, Southern Biotec, Alabama, USA). Staining steps were performed at 4 °C for 30 min. PBMC were then analysed by flow cytometry; only ~28% of the cells within the PBMC expressed BoLA-DR on the cell surface; however, of these, ~90% had internalised fOVA (Supplementary Figure 1b). Having optimised the conditions for loading the cells with exogenous proteins ovalbumin was used in large scale cultures to generate the samples for pBoLA-DR elution analysis.

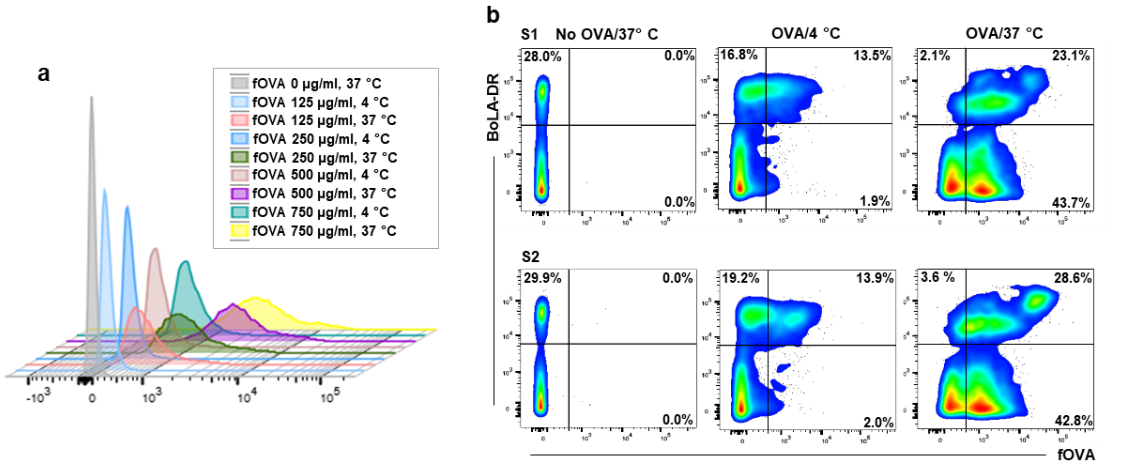


**Supplementary Figure 1 - Optimization of cell incubation conditions for exogenous antigen presentation assay.** Detection of fOVA+ cells was performed by flow cytometry. a) fOVA internalization by *T. annulata*-infected cell line; analysis of the percentage of T. annulata infected cells fluorescent after incubation with different concentrations of fOVA at either 4°C or 37°C. b) fOVA internalization and BoLA-DR quantification in PBMCs. S1: test sample 1. S1: test sample 2. 4°C samples were used as passive-internalization controls.

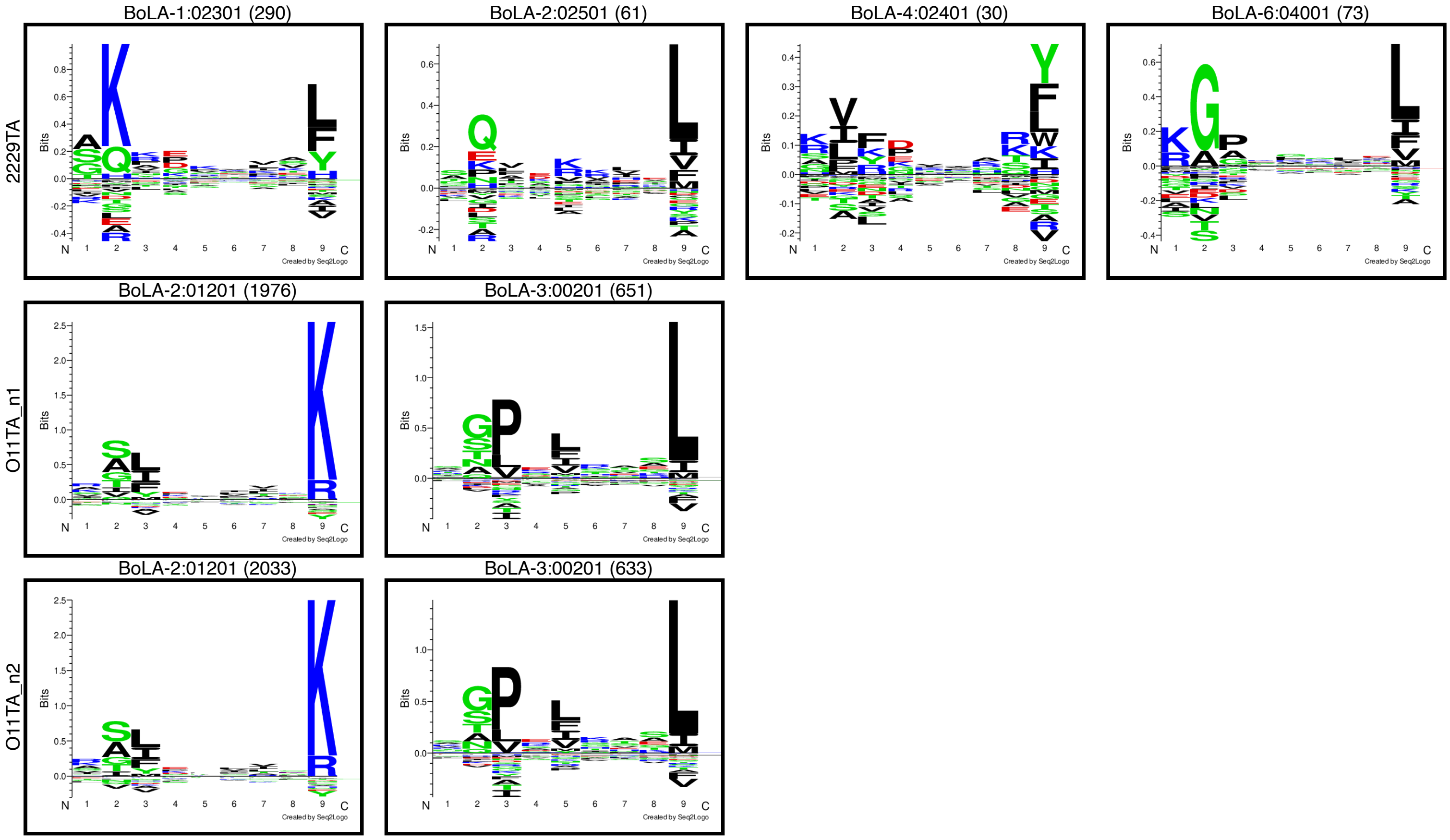


**Supplementary Figure 2 - MHCI motifs deconvoluted from peptides from samples 2229TA, O11TA_n1 and O11TA_n2.** Peptides of length 8-13 from the samples were deconvoluted and their binding potential to the MHCI alleles in the respective haplotypes was predicted using NetMHCpan-4.1. Logos were generated from predicted binding cores of peptides with rank lower than 5%. Numbers in parentheses indicate the number of peptides assigned to each allele.

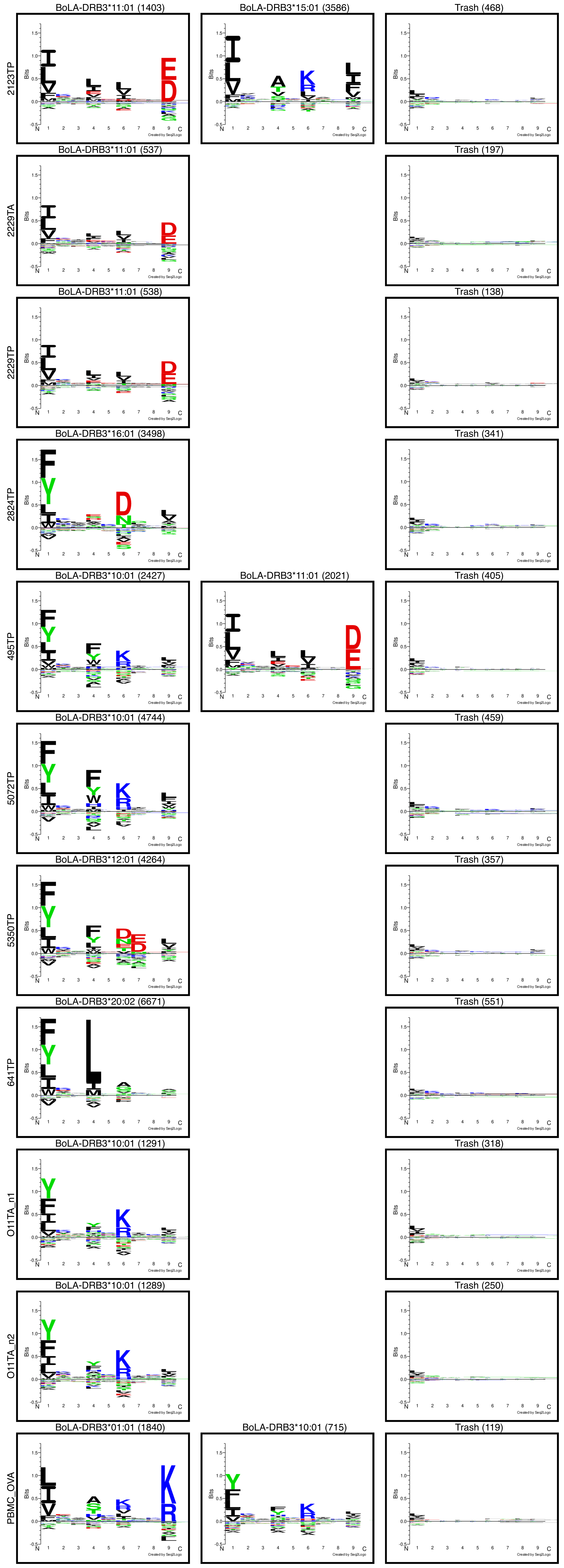


**Supplementary Figure 3 - MHCII binding core logos for deconvolution solutions for samples in BoLA training data.** Each logo is generated from binding core alignments of peptides assigned to the given allele with a prediction rank score <20%, as predicted by the BoLA model with context encoding. Poorly ranking peptides of each sample are assigned to a trash cluster. Numbers in parentheses indicate the number of peptides assigned to each cluster. All logos have a constant y-axis range.

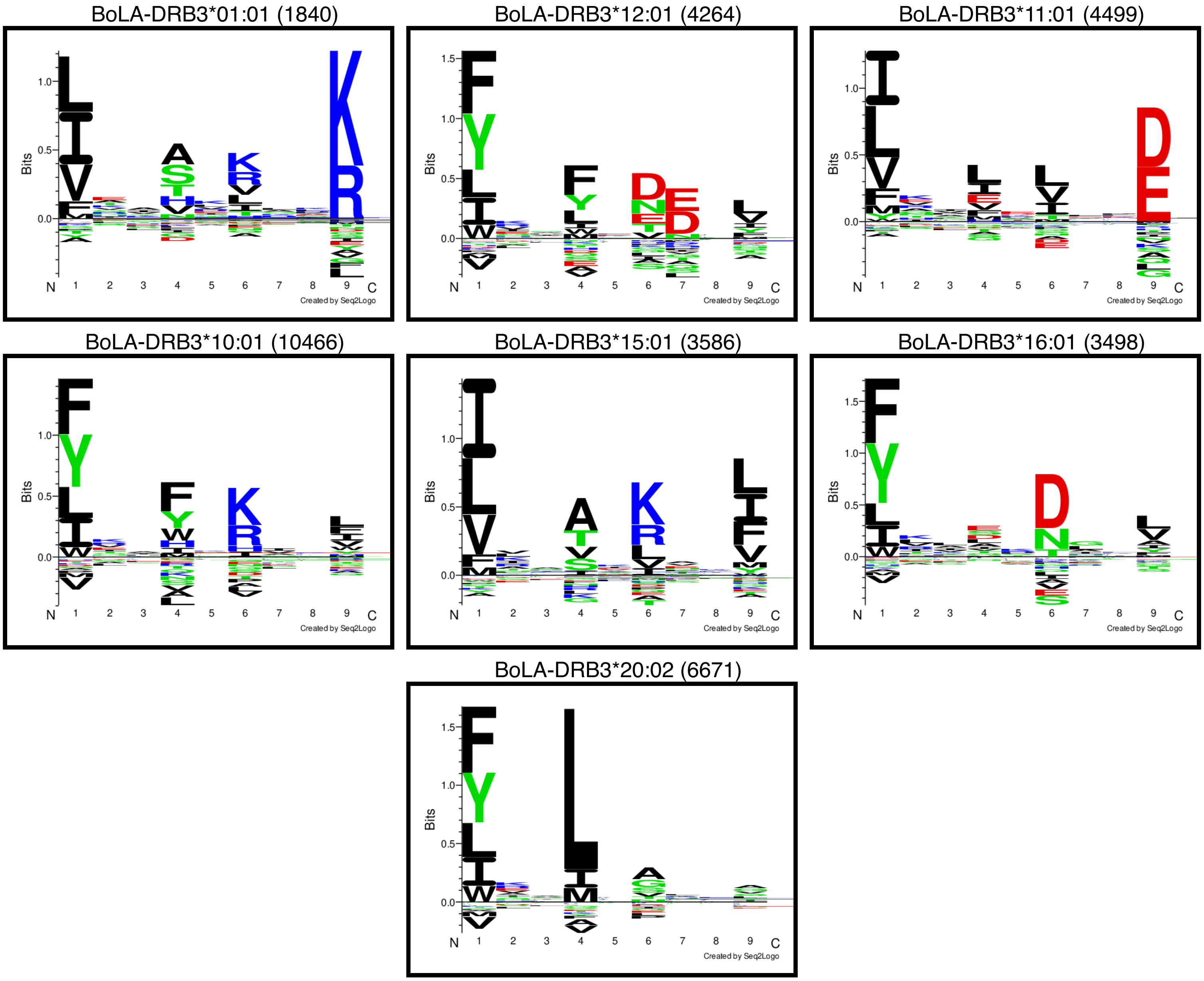


**Supplementary Figure 4 - MHCII binding core motifs for all alleles in BoLA training data, combining allele data across samples.** Each logo is generated from binding core alignments of peptides assigned to the given allele with a prediction rank score <20%, as predicted by the BoLA model with context encoding. Numbers in parentheses indicate the number of peptides assigned to each allele.

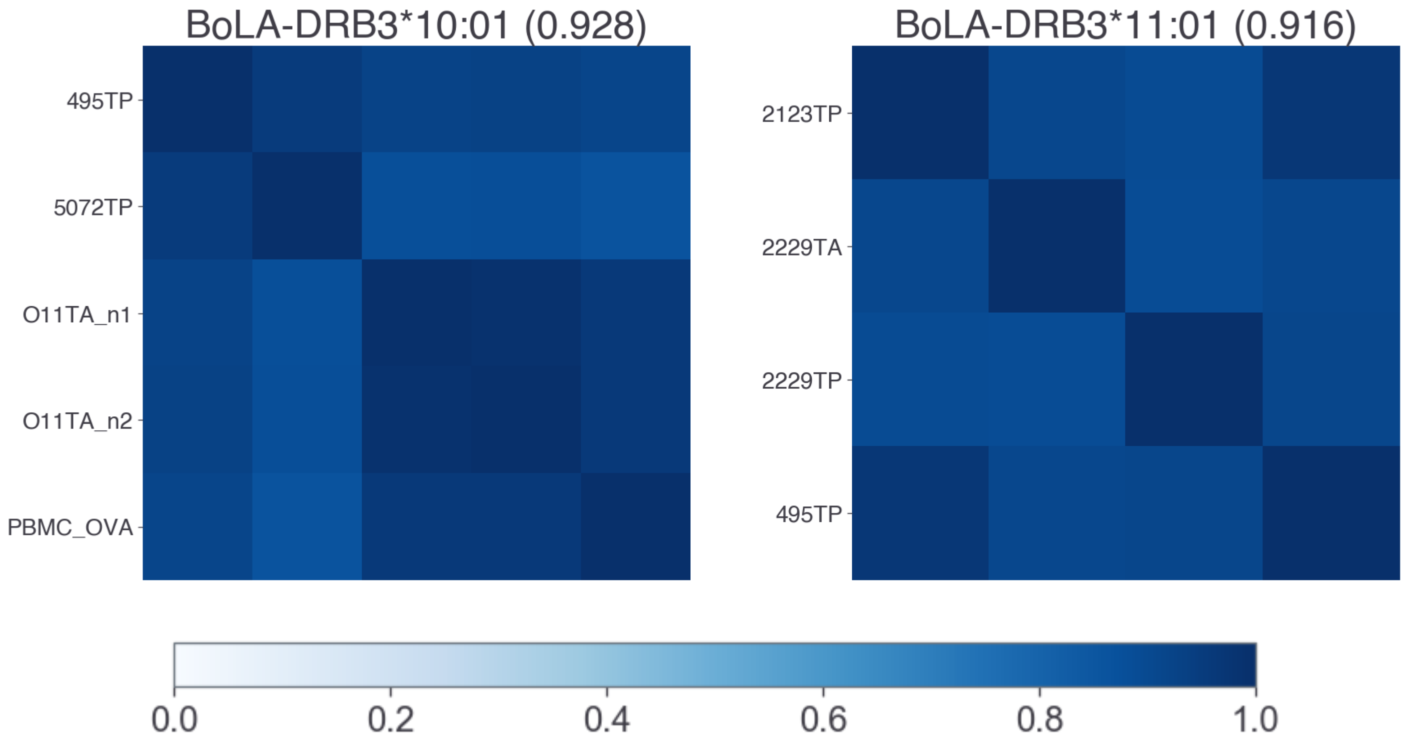


**Supplementary Figure 5 - Deconvolution consistency heatmaps for alleles observed across multiple samples.** The shading of the heatmaps indicates the Pearson correlation comparing allele logos across different samples. The value in parentheses is the consistency score, which is the average correlation in the matrix.

**Supplementary Table 1 – PPV values for deconvolution of BoLA data as predicted by the BoLA model with context encoding.**

| *Sample* | *MHC* | *#Ligands* | *#Ligands <Rank 20* | *#Negatives* | *PPV* |
|---|---|---|---|---|---|
| *2123TP* | BoLA-DRB3*11:01 | 1417 | 1403 | 2597 | 0.765 |
| *2123TP* | BoLA-DRB3*15:01 | 4040 | 3586 | 43289 | 0.835 |
| *2229TA* | BoLA-DRB3*11:01 | 734 | 537 | 7097 | 0.775 |
| *2229TP* | BoLA-DRB3*11:01 | 676 | 538 | 6204 | 0.794 |
| *2824TP* | BoLA-DRB3*16:01 | 3839 | 3498 | 34914 | 0.849 |
| *495TP* | BoLA-DRB3*10:01 | 2805 | 2427 | 38998 | 0.788 |
| *495TP* | BoLA-DRB3*11:01 | 2048 | 2021 | 2851 | 0.787 |
| *5072TP* | BoLA-DRB3*10:01 | 5203 | 4744 | 40940 | 0.851 |
| *5350TP* | BoLA-DRB3*12:01 | 4621 | 4264 | 35866 | 0.864 |
| *641TP* | BoLA-DRB3*20:02 | 7222 | 6671 | 66059 | 0.834 |
| *O11TA_n1* | BoLA-DRB3*10:01 | 1609 | 1291 | 19055 | 0.796 |
| *O11TA_n2* | BoLA-DRB3*10:01 | 1539 | 1289 | 17707 | 0.82 |
| *PBMC_OVA* | BoLA-DRB3*01:01 | 1843 | 1840 | 1409 | 0.868 |
| *PBMC_OVA* | BoLA-DRB3*10:01 | 831 | 715 | 26231 | 0.751 |

**Supplementary Table 2 – *T. Parva* CD4 epitopes in the CD4 benchmark.**

| *UID* | *MHC* | *Peptide* |
|---|---|---|
| *TpMuguga_01g00726* | BoLA-DRB3*10:01 | GFLGDNMIDKSDKMPWYK |
| *TpMuguga_03g00655* | BoLA-DRB3*10:01 | GRVSNYVTYAKKLLSNGI |
| *TpMuguga_03g00861* | BoLA-DRB3*10:01 | QEILYYKWEKHGFVKETY |
| *TpMuguga_03g00861* | BoLA-DRB3*10:01 | PLSGYHVRYVNYGKVIMW |
| *TpMuguga_04g00916* | BoLA-DRB3*10:01 | HPGIQYVPYQTLQI |
| *TpMuguga_04g00916* | BoLA-DRB3*10:01 | QPTQIEQSGTYQHYGPPVFP |
| *TpMuguga_02g00244* | BoLA-DRB3*10:01 | NEEYAAFYKNLTNDWEDH |
| *TpMuguga_02g00895* | BoLA-DRB3*11:01 | YDGEKVWSLEVGGDYA |
| *TpMuguga_02g00895* | BoLA-DRB3*11:01 | EKTIEITFIGGEKEIY |
| *TpMuguga_01g00726* | BoLA-DRB3*11:01 | ITGTSQADVAMLVVPAES |
| *TpMuguga_01g00726* | BoLA-DRB3*11:01 | GKILVEALDLMEPPKRPV |
| *TpMuguga_02g00123* | BoLA-DRB3*11:01 | ELTLEGIKQFYILIDKEY |
| *TpMuguga_04g00437* | BoLA-DRB3*11:01 | KSFDDLTTVELAPEPKAS |
| *TpMuguga_04g00916* | BoLA-DRB3*16:01 | IEMTEKEYKIIVDSRFKIKF |
| *TpMuguga_04g00683* | BoLA-DRB3*16:01 | QIEVTFNIDTNGILSVTA |
| *TpMuguga_04g00752* | BoLA-DRB3*16:01 | NCGRGVFMAAHNNRTYCG |
| *TpMuguga_03g00861* | BoLA-DRB3*16:01 | QEILYYKWEKHGFVKETY |
| *TpMuguga_03g00861* | BoLA-DRB3*16:01 | FNKFDMLHDGVYYSSPVP |
| *TpMuguga_03g00861* | BoLA-DRB3*16:01 | CKANNPVVYIKAGDKTVW |
| *TpMuguga_01g01077* | BoLA-DRB3*16:01 | KYFAIDIDGTFHIKD |
| *TpMuguga_01g01077* | BoLA-DRB3*16:01 | ATPPKYFAIDIDGTFHI |
| *TpMuguga_01g01074* | BoLA-DRB3*16:01 | KYFAIDIDGTFFIKD |
| *TpMuguga_01g01074* | BoLA-DRB3*16:01 | EKLPKYFAIDIDGTFFI |
| *TpMuguga_01g01074* | BoLA-DRB3*16:01 | NIAAFKRLQDAGVLPFF |
| *TpMuguga_02g00895* | BoLA-DRB3*16:01 | YDGEKVWSLEVGGDYAVKV |

**Supplementary File 6 - Applying NetBoLAIIpan to the rational screening of vaccine candidates in complex parasites**

Ticks are a major pathogen of livestock, causing disease both directly and as vectors of several diseases such as ehrlichiosis, anaplasmosis, babesiosis and theileriosis. The most economically significant tick globally is *R. microplus;* in Brazil alone, the annual losses due to *R. microplus* infestation is estimated to be USD 3.2 billion(1). Attempts to develop novel vaccines against ticks often target salivary proteins as candidate antigens. However, our group's recent work has highlighted the complexity of the sialotranscriptome of *R. microplus*, with ~3600 CDS identified as being potentially secreted proteins(2). Rationally selecting candidate antigens from this large number of proteins is a major obstacle to vaccine development. Three families of tick salivary proteins that have been used as targets for vaccines are the lipocalins, Kunitz domain-containing protease inhibitors and cement proteins. Members from each of these protein families have been used in experimental vaccines against ticks and shown some degree of protective efficacy(3–6). These results indicate that proteins from these families are promising candidates for vaccine development in mono- or multi-antigenic configurations for *R. microplus*.

Analysis of the *R. microplus* sialotranscriptome(1) identified a total of 294 lipocalins, 201 Kunitz domain-containing protease inhibitors, and 49 cement family members. Deciding which of these proteins to take forward into immunogenicity/efficacy trials requires some parameters to select particular proteins or combinations of proteins preferentially. To evaluate if the NetBoLAIIpan algorithm could contribute to the rational preliminary selection, we incorporated it as part of a pipeline to analyse proteins from these three families. First, the sialotranscriptome was filtered to remove all sequences that had >3% unresolved amino acids. Then the ten proteins from each of these three families that demonstrated the highest level of expression (by transcript number identified in nymphs and adult ticks fed on Holstein-Friesian cattle) were selected. These 30 proteins were then analysed for peptides presented by the seven BoLA-DR molecules included in this study (using a threshold of predicted rank binding of <1%). Also included in this analysis was the *R. microplus* midgut glycoprotein Bm86 protein – this is the only tick antigen that has been used in a commercially available tick vaccine(7). The output of this data is presented in Supplementary Table 3.

**Supplementary Table 3 - BoLA-DRB3 ligand prediction in salivary proteins from *R. microplus* tick.**

| | | DRB3 alleles/#strong binders | | | | | | | |
|---|---|---|---|---|---|---|---|---|---|
| Protein Family | Protein ID | *0101 | *1001 | *1101 | *1201 | *1501 | *1601 | *2002 | No. of BoLA-DR molecules with predicted strong binders |
| Lipocalins | Rm-nr-78292 | 0 | 1 | 0 | 0 | 1 | 3 | 0 | 3 |
| | Rm-nr-78501 | 2 | 0 | 0 | 1 | 0 | 2 | 1 | 4 |
| | Rm-tb2-53013 | 6 | 6 | 0 | 5 | 2 | 4 | 2 | 6 |
| | Rm-nr-74000 | 0 | 1 | 0 | 3 | 0 | 3 | 0 | 3 |
| | Rm-nr-78500 | 2 | 0 | 1 | 2 | 0 | 5 | 1 | 5 |
| | Rm-nr-78498 | 1 | 0 | 1 | 2 | 0 | 2 | 1 | 5 |
| | Rm-nr-73997 | 2 | 0 | 0 | 0 | 0 | 1 | 0 | 2 |
| | Rm-tb2-50143 | 1 | 1 | 5 | 1 | 0 | 1 | 0 | 5 |
| | Rm-tb2-8240 | 5 | 2 | 4 | 1 | 0 | 0 | 6 | 5 |
| | Bm-tb2-48009 | 4 | 2 | 3 | 1 | 0 | 2 | 3 | 6 |
| KU | Rm-nr-115992 | 0 | 3 | 1 | 0 | 1 | 0 | 0 | 3 |
| | Rm-nr-95340 | 0 | 2 | 0 | 0 | 0 | 0 | 0 | 1 |
| | Rm-nr-113888 | 0 | 0 | 0 | 1 | 0 | 0 | 0 | 1 |
| | Rm-tb2-95338 | 0 | 1 | 0 | 0 | 0 | 0 | 0 | 1 |
| | Rm-nr-32664 | 0 | 0 | 0 | 1 | 0 | 3 | 0 | 3 |
| | Rm-nr-95339 | 0 | 1 | 0 | 0 | 0 | 0 | 0 | 1 |
| | Rm-tb2-16567 | 0 | 1 | 0 | 0 | 1 | 1 | 0 | 3 |
| | Rm-nr-105260 | 0 | 1 | 0 | 0 | 0 | 1 | 0 | 2 |
| | Rm-nr-41891 | 0 | 0 | 0 | 0 | 0 | 0 | 0 | 0 |
| | Rm-tb2-10954 | 0 | 0 | 0 | 0 | 0 | 0 | 0 | 0 |
| Cement | Rm-nr-4894 | 0 | 0 | 2 | 5 | 0 | 1 | 5 | 4 |
| | Rm-nr-8830 | 0 | 0 | 0 | 4 | 0 | 0 | 3 | 2 |
| | Rm-nr-2649 | 0 | 0 | 0 | 0 | 0 | 0 | 6 | 1 |
| | Rm-tb2-8829 | 0 | 0 | 2 | 4 | 2 | 1 | 5 | 5 |
| | Rm-nr-34342 | 3 | 1 | 0 | 0 | 0 | 2 | 6 | 4 |
| | Rm-nr-8834 | 3 | 2 | 1 | 0 | 0 | 0 | 8 | 4 |
| | Rm-tb2-8267 | 0 | 2 | 0 | 0 | 0 | 0 | 0 | 1 |
| | Rm-nr-9404 | 0 | 0 | 0 | 0 | 0 | 0 | 1 | 1 |
| | Rm-nr-2167 | 0 | 0 | 0 | 0 | 0 | 0 | 7 | 1 |
| | Rm-nr-2175 | 0 | 0 | 0 | 0 | 0 | 0 | 5 | 1 |
| Bm86 | P20736 | 1 | 0 | 1 | 1 | 5 | 1 | 1 | 6 |

| | | | | | | | |
|---|---|---|---|---|---|---|---|
| No. of proteins containing predicted strong binders (n/30) | 11 | 15 | 10 | 14 | 6 | 16 | 16 |

Importantly, no single protein was found to contain peptides that were predicted to be strong binders for all of seven BoLA-DR molecules. In the lipocalin family, two proteins (Rm-tb_53013 and Rm-tb2-48009) had predicted binders to 6/7 of the BoLA-DR molecules; in the cement family the maximum number of BoLA-DR molecules covered by any single protein was only five (Rm-tb2-8829), and in the Kunitz domain-containing protease inhibitor family no protein had binders for more than three of the BoLAD-DR molecules. The Bm86 protein contained predicted binders for 6/7 of the BoLA-DR molecules. On average, a predicted binder for any single BoLA-DR molecule was only found in ~1/3 of proteins (12.6/31 – a range of 7-17/31 proteins), while on average each protein only contained predicted binders for ~3 BoLA-DR molecules. Such observations suggest that to ensure the inclusion of peptides that are predicted to be able to be presented by a range of BoLA-DR molecules requires careful selection of complementary candidate antigens for inclusion in a vaccine. Before the availability of the NetBoLAIIpan algorithm, such *in silico* analysis was not feasible and candidate antigen selection (which has often been based solely on parameters such as transcription levels and antibody immunogenicity) may have led to less than optimal antigen combinations being taken forward for assessment in expensive and laborious *in vivo* animal experiments.

In summary, this small-scale exemplar has demonstrated how the application of NetBoLAIIpan for preliminary *in silico* prediction of BoLA-DR binders could have a profound influence on the selection of candidate antigens by providing predictions of which antigen combinations should facilitate stimulation of CD4 T cells in animals expressing a range of BoLA-DR molecules. Further data expanding the repertoire to incorporate more of the high-frequency DRB3 alleles and comparable data for the more complex BoLA-DQ system would enable more comprehensive analyses. Integrating such immuno-informatic approaches early could facilitate a rapid, cheap and more rational way of selecting candidate antigens from highly complex pathogens (which often express hundreds or thousands of proteins) for subsequent *in vivo* assessment; this could have significant implications and benefits for the development of vaccines against *R. microplus* and other eukaryotic parasites.
